# DNA damage and senescence in the aging and Alzheimer’s disease cortex are not uniformly distributed

**DOI:** 10.1101/2024.04.28.591529

**Authors:** Gnanesh Gutta, Jay Mehta, Rody Kingston, Jiaan Xie, Eliana Brenner, Fulin Ma, Karl Herrup

## Abstract

Alzheimer’s disease (AD) is a neurodegenerative illness with a typical age of onset exceeding 65 years of age. The age-dependency of the condition led us to track the appearance of DNA damage in the frontal cortex of individuals who died with a diagnosis of AD. The focus on DNA damage was motivated by evidence that increasing levels of irreparable DNA damage are a major driver of the aging process. The connection between aging and the loss of genomic integrity is compelling because DNA damage has also been identified as a possible cause of cellular senescence. The number of senescent cells has been reported to increase with age, and their senescence-associated secreted products are likely contributing factors to age-related illnesses. We tracked DNA damage with 53BP1 and cellular senescence with p16 immunostaining of human post-mortem brain samples. We found that DNA damage is significantly increased in the BA9 region of the AD cortex when compared to the same region of unaffected controls (UC). In the AD but not UC cases, the density of cells with DNA damage increased with distance from the pia mater up to approximately layer V then decreased in deeper areas. This pattern of DNA damage was overlaid with the pattern of cellular senescence, which also increased with cortical depth. On a cell-by-cell basis, we found that the intensity of the two markers was tightly linked in the AD, but not the UC brain. To test whether DNA damage was a causal factor in the emergence of the senescence program, we used etoposide treatment to damage the DNA of cultured mouse primary neurons. While DNA damage increased after treatment, after 24 hours no change in the expression of senescence-associated markers was observed. Our work suggests that DNA damage and cellular senescence are both increased in the AD brain and increasingly coupled. We propose that in vivo the relationship between the two age-related processes is more complex than previously thought.

## 1. INTRODUCTION

The most prevalent neurodegenerative diseases – Alzheimer’s, Parkinson’s, amyotrophic lateral sclerosis, Huntington’s, and others – all share age as a common risk factor. A particularly stunning example of this age-dependence is the late-life dementing illness known as Alzheimer’s disease (AD). Before the age of 65, AD is uncommon 0.1% prevalence - ^1^. Seen in the context of the average life expectancy (approximately 79 years in the United States) people with Alzheimer’s Disease would be expected to have lived over 80% of their lives with a brain, capable of healthy neural network activity and no overt evidence of AD symptoms. The biological puzzle that these simple observations present is that somehow, after years of healthy mental functioning, advancing age somehow increases the risk of a degenerative process that results in the characteristic set of symptoms we recognize as Alzheimer’s Disease (AD) – short-term memory loss, behavioral symptoms, depression, apathy, and ultimately loss of basic homeostasis and death.

The cell and molecular relationships between aging and AD are difficult to uncover in large part because the biology of aging is incompletely understood. Metabolic imbalances altering flux through the insulin signaling pathway contribute to aging ^2^ as does the loss of basic cellular functions such as mitochondrial integrity ^3^ or autophagy ^4^. While many factors are suspected to contribute to the process of aging, the existing data largely support the hypothesis that the loss of genomic integrity is the major driver ^5-10^. First, there is a tight correlation between age and the accumulation of unrepaired DNA damage ^11-13^. In addition, mutations in proteins needed for DNA repair, such as ataxia-telangiectasia mutated (ATM), WRN (the gene mutated in Werner syndrome), and others lead to conditions whose phenotypes include premature aging ^3, 14, 15^. This latter observation represents a compelling argument that DNA damage is not just correlated with age but may help to drive it. This would seem logical since without an intact genome proper cellular function is all but impossible. This theory of aging has relevance to Alzheimer’s disease. While DNA damage increases with normal aging, it increases far more rapidly in the brains of persons with AD. This connection has been recognized for over 30 years ^16, 17^, and its significance has been validated with modern molecular methods ^18-22^.

DNA damage is a constant in the life of any cell with tens of thousands of lesions estimated to alter the genome of every cell, every day ^23^. Compounding the problem for the nervous system is that relatively modest neuronal activity can itself induce DNA damage ^24, 25^. To deal with these problems, cells have a wide array of DNA repair proteins that quickly and efficiently repair lesions in the double helix. No repair process is perfect, however, and with time irreparable damage accumulates. In addition, for reasons that are poorly understood, the levels of many DNA repair proteins decrease with age ^26, 27^, which likely contributes to the increased levels of DNA damage with age.

In addition to contributing to the process of aging, many DNA lesions destroy the ability of the cell to maintain a differentiated, non-dividing state, thus opening the door to malignant transformation and cancer. To guard against this, our cells are programmed to respond to excessive DNA damage by entering a cellular state known as senescence. Senescent cells have many intriguing properties, but among the most important of these is a total cell cycle arrest ^28, 29^. As with DNA damage, the number of cells with evidence of senescence increases with age. This compounds the problems associated with DNA damage alone as senescent cells are known to release pro-inflammatory factors collectively known as the Senescence-Associated Secretory Phenotype (SASP). SASP acts as a signal of potential danger to surrounding cells, similar to an innate immune response. It can also induce senescence in neighboring cells, enhancing aging in the region. While the signaling by SASP initially protects the tissue, the resulting chronic inflammatory state is maladaptive by making tissues more vulnerable to age-related disorders, such as Alzheimer’s Disease (AD). The density of senescent cells increases in the AD forebrain, particularly in cells containing neurofibrillary tangles (Saez-Atienzar & Masliah, 2020).

The literature in this area has an important gap, however. Most studies of DNA damage, senescence and brain aging report results either at the level of the single cell or use solution-based methods such as western blots that would mask any regional heterogeneity. Regional variation has been observed in young animals ^24^, but its extent is rarely examined in older experimental animals or in humans. To begin to fill this gap, we have asked whether the levels of DNA damage or senescence that accumulate with age and Alzheimer’s disease differ with depth in the human cerebral cortex. We report here that such variations are found and discuss the implications of these findings for aging, senescence, and Alzheimer’s disease.

## 2. Materials and Methods

### 2.1. Human samples

The human samples we used in experiments were obtained from the National Institutes of Health NeuroBioBank at the University of Maryland, Baltimore, MD, the Human Brain and Spinal Fluid Resource Center, and the Mount Sinai/James J. Peters VA Medical Center, National Institutes of Health Brain and Tissue Repository, with approvals from Tissue Access Committee of NeuroBioBank. The samples were all from the Brodmann area 9 region (BA9) of the left frontal cortex. Age at death ranged from 76 to 82 years. Clinical diagnosis of patients was done based on Braak staging with Braak stages 0-II being categorized as UC, and Braak Stages V-VI being categorized as AD. The presence of amyloid pathology was confirmed by immunostaining for amyloid plaques using the 6E10 monoclonal antibody (Fig. 1). We analyzed 4 cases from persons who had been diagnosed with AD. Although genotyping was not performed, the relatively advanced age of death makes it highly likely that these cases all suffered from sporadic Alzheimer’s disease. Both AD and UC cases had a postmortem interval of less than 10 h. Brain tissues were flash frozen without fixation and were stored at -80°C before they were cryosectioned at 13 μm and mounted on SuperFrostPlus glass slides.

**Figure 1.**
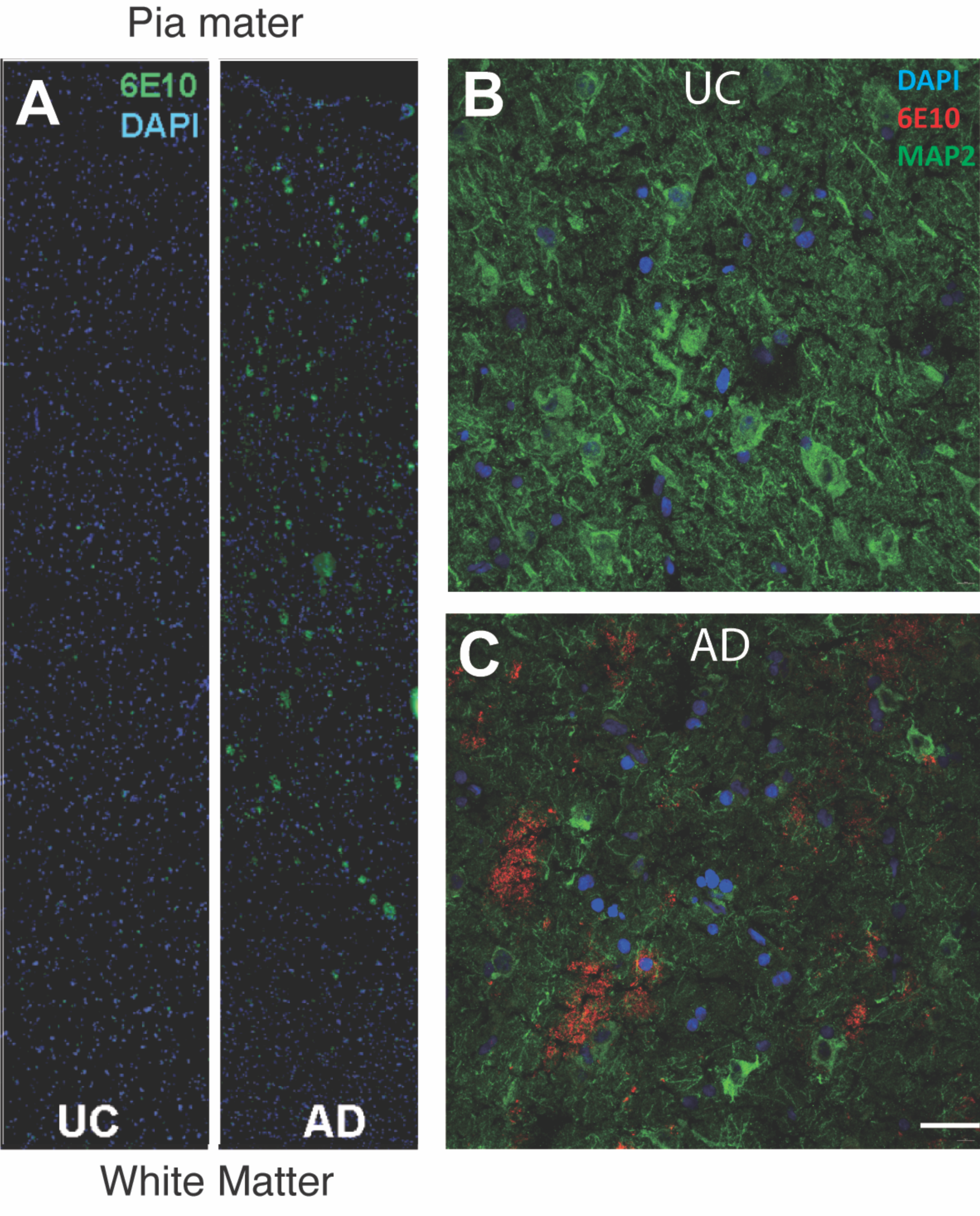
Representative images of amyloid deposit density in AD and UC brain. **A**) Radial fields are shown spanning the entire distance from the pia mater (top) to the white matter (bottom) in the sampled regions of BA9. The left panel illustrates the 6E10 immunostaining pattern in UC material (Braak 0-II). Deposits of amyloid are small and relatively sparse. The right panel illustrates a comparable image from an AD case. Note the increased density of 6E10-positive deposits found largely in the upper third of the cortex. **B**) A higher magnification of a similar section from a UC case immunostained with the indicated antibodies. **C**) A comparable field from an AD case showing the dense nature of the amyloid deposits (red).

### 2.2. Primary Culture

The surface of each cover slip was coated with 0.5 mg/ml poly-L-lysine diluted in boric acid buffer (Sigma). Embryos were harvested from pregnant, wildtype C57BL/6J mice (IMSR_JAX:000664) on embryonic day 16.5 (E16.5) and decapitated. The cerebral cortices were dissected free of meninges and stored at 4°C in PBS with 1 mg/ml glucose. Cortices were then minced with small forceps and incubated at 37°C in 0.25% trypsin-EDTA (Sigma) for 10 min at 37°C. The trypsin was inactivated with 10% FBS in DMEM after which the tissue was rinsed with Neurobasal medium. The tissue was gently aspirated to create a single cell suspension and the constituent cells plated in 24-well plates on the coated coverslips at 30,000 cells/well. The cultures were maintained at 37°C in a humidified atmosphere of 5% CO_2_ in Neurobasal medium supplemented with 2% B27 (Invitrogen), 1% Glutamax (Invitrogen), and 10,000 U/ml penicillin/streptomycin (Invitrogen). Every 5 d, the culture medium was refreshed by replacing half of the old medium with fresh.

### 2.3. Immunocytochemistry

#### 2.3.1. Tissue sections

Sections were warmed to room temperature, then rinsed in PBS three times for 5 min each time. Non-specific binding was blocked by incubating the sections at room temperature for 1 hr in 0.3% Triton X-100 in PBS (PBST) with 10% donkey serum. After blocking, the tissues were incubated overnight at 4°C with primary antibody in 10% donkey serum in PBST at dilutions listed in Table I. The following day, tissues were rinsed in PBS three times for 5 min each time and incubated at room temperature for 1 hr with secondary antibodies at a 1:500 dilution in 10% donkey serum in PBST. The tissues were then rinsed in PBS three times for 5 min each time and stained with DAPI at a 1:100 dilution in PBST for 10 min at room temperature. After two final rinses in PBS the stained sections were mounted with Hydromount. Sections were viewed either on a fluorescence microscope (Nikon Eclipse Ni-U) or on a confocal microscope (Nikon Eclipse Ti2).

**TABLE I.** Antibodies used.

| **TABLE I** | **Antibodies used** |  |  |  |
| --- | --- | --- | --- | --- |
| **Target antigen** | **Manufacturer** | **Catalog #** | **RRID #** | **Dilution** |
| MAP2 | Abcam | ab5392 | AB_2138153 | 1:2500 |
| 53BP1 | Abcam | ab36823 | AB_722497 | 1:500 |
| CDKN2A/p16INK4a | Abcam | ab54210 | AB_881819 | 1:500 |
| γ-H2AX | Abcam | ab26350 | AB_470861 | 1:500 |
| p27<sup>KIP1</sup> [Y236] | Abcam | ab32034 | AB_2244732 | 1:400 |

Whole slide images (WSI) were generated using a Zeiss Axioscan.Z1 and Colibri v7 LED excitation source (Carl Zeiss, Oberkochen, Germany), equipped with a cooled 16-bit CMOS monochrome camera (Hamamatsu Photonics, Japan) for imaging. Acquisition resolution was 0.325 µm/pixel (via a 20x, 0.8 N.A. objective). WSI acquisition parameters (focus, exposure times, excitation intensity) were adjusted specifically for optimization of the dynamic range for each fluorophore used and kept constant across all imaging sessions.

#### 2.3.2. Primary cultures

Cells on coverslips were rinsed in PBS then fixed in 4% paraformaldehyde for 15 minutes then washed with PBS three times (5 min each) and blocked for 1 hr with 5% donkey serum in PBST. The coverslips were then incubated overnight at 4°C with primary antibodies at the dilutions shown in Table I in a solution containing 5% donkey serum in PBST. The following day, cells were washed in PBS (3 x 5 min) and incubated for one hour at room temperature with secondary antibodies diluted 1:500 with PBST containing 5% donkey serum. Coverslips were washed with PBS (3 x 5 min) and incubated with DAPI diluted 1:50 in PBST for 10 min at room temperature. After two additional rinses in PBS the coverslips were mounted with Hydromount.

For each biological repeat, we analyzed at least two coverslips, and for each coverslip 3 regions of interest (ROI) were selected and imaged. The ROIs were selected at 4x magnification using the MAP2 channel to ensure adequate neuronal density. Then, an image was captured sequentially by the monochrome camera in all fluorescent channels to generate a merged image for the quantification described. Fluorescent imaging was done on a Nikon Eclipse Ni-U microscope with a 20× objective and equipped with DAPI (400-418 nm), FITC (450-490 nm), TRITC (554-570 nm), and Cy5 (640-670 nm) filter sets controlled by NIS-Elements AR software.

#### 2.3.3. Image analysis

For quantification of the images, QuPath, an open-source software for digital pathology and whole slide image analysis, was used ^30^. The largest continuous area of the gray matter was identified, and evenly spaced vertical lines were drawn from the white matter to the pia defining vertical columns within the identified area. Using the coordinates displayed by QuPath, the two endpoints plus five equidistant points were made on each line and connected using the draw annotation to define six radial zones (Figure 1). These zones are similar, but not identical, to the six cortical layers as defined by histological criteria. Once the six zones were annotated, the positive cell detection tool was used to identify cells of interest. Parameters were then adjusted to ensure the software was accurately detecting DAPI nuclear staining as confirmed by spot manual counts. The nuclear area was artificially expanded by 5 μm following the shape of the nucleus to create an artificial “pseudoplasm”. To detect DNA damage in the cells, object classification was used to quantify 53BP1 positive cells among the DAPI-positive nuclei that were previously detected. The appropriate parameters were set and confirmed by spot manual counts throughout the image to determine a threshold level that accurately quantified the 53BP1 staining. Above the threshold was considered positive for 53BP1 and below the threshold was ignored. Once the measurements were set, data was exported as a .csv file then analyzed using Excel and Prism software.

### 2.4. Statistical analysis

All statistical analysis was performed using GraphPad Prism software. Data is presented for each group as mean ± SEM. The statistical significance of changes between different groups was evaluated using unpaired t tests with p values <0.05 being considered significant. To compare across cortical layers, we used ANOVA with multiple comparisons.

## 3. Results

### 3.1. DNA damage and senescence are present in both the unaCected and Alzheimer’s disease brains

We verified the presence of cells with persistent DNA damage in the cortical gray matter of cases that had come to autopsy with a diagnosis of Alzheimer’s disease and that we subsequently verified to have the expected density of amyloid plaques. DNA damage was also present in age-matched unaffected control (UC) cases for which there was no history of cognitive impairment. To detect DNA damage, we immunostained cryostat sections of Brodmann Area 9 (BA9) with both 53BP1 and *γ*-H2AX (Fig. 2). We found a very high degree of overlap of the two DNA damage markers in both UC (top panels) and AD (bottom panels). Both antibodies stained a small percentage of the total number of cells in radial locations ranging from the pia to the white matter. The staining was predominantly nuclear in location and could be found in both large and small nuclei. As reported by others, the density of nuclei with evidence of DNA damage was clearly higher in the AD cases than in the unaffected controls (UC) and this was true at all cortical depths (compare Figs. 3B and D with Figs. 3A and C). The senescence marker, p16^INK4a^ was also readily apparent in the cells in the same areas (Fig. 3). The particulars of its cellular localization and the implications for cellular senescence are discussed in more detail below.

**Figure 2.**
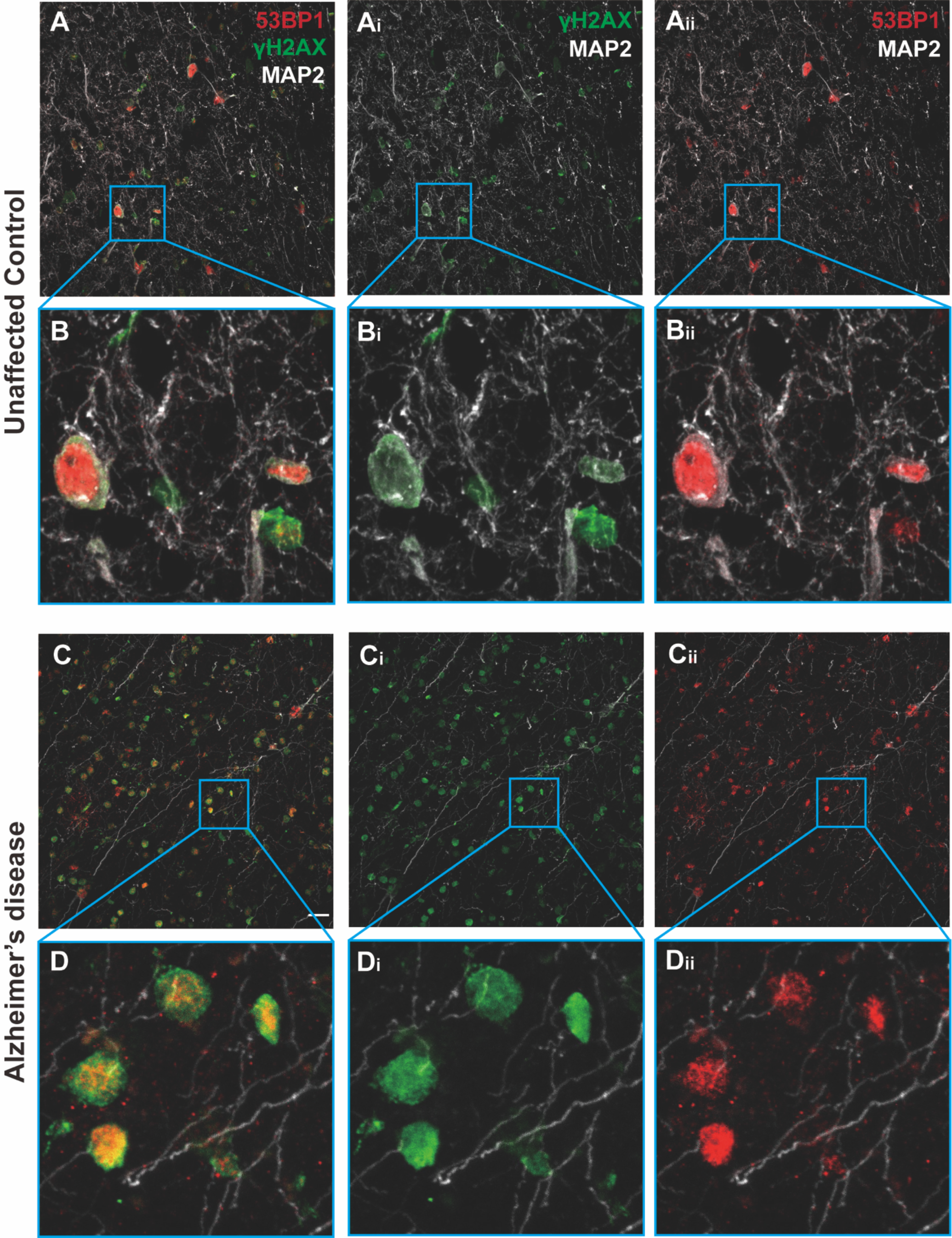
Typical images from the middle layers of BA9 cortex immunostained for two DNA damage markers, 53BP1 (red) and *γ*-H2AX (green) plus MAP2 (white). Note the near total overlap in the nuclei of immunostained cells **A**) Low magnification view of a region from an unaffected control (UC). **Ai**) The same field showing only *γ*-H2AX and MAP2 staining. **Aii**) The same field showing only 53BP1 and MAP2 staining. **B**, **Bi**, **Bii**) Higher magnification of the areas indicated by the blue boxes. **C**, **Ci**, **Cii**) Images comparable to those shown A, but showing a region from an Alzheimer’s disease brain. **D**, **Di**, **Dii**) Images comparable to those shown B, but showing a region from an Alzheimer’s disease brain.

**Figure 3.**
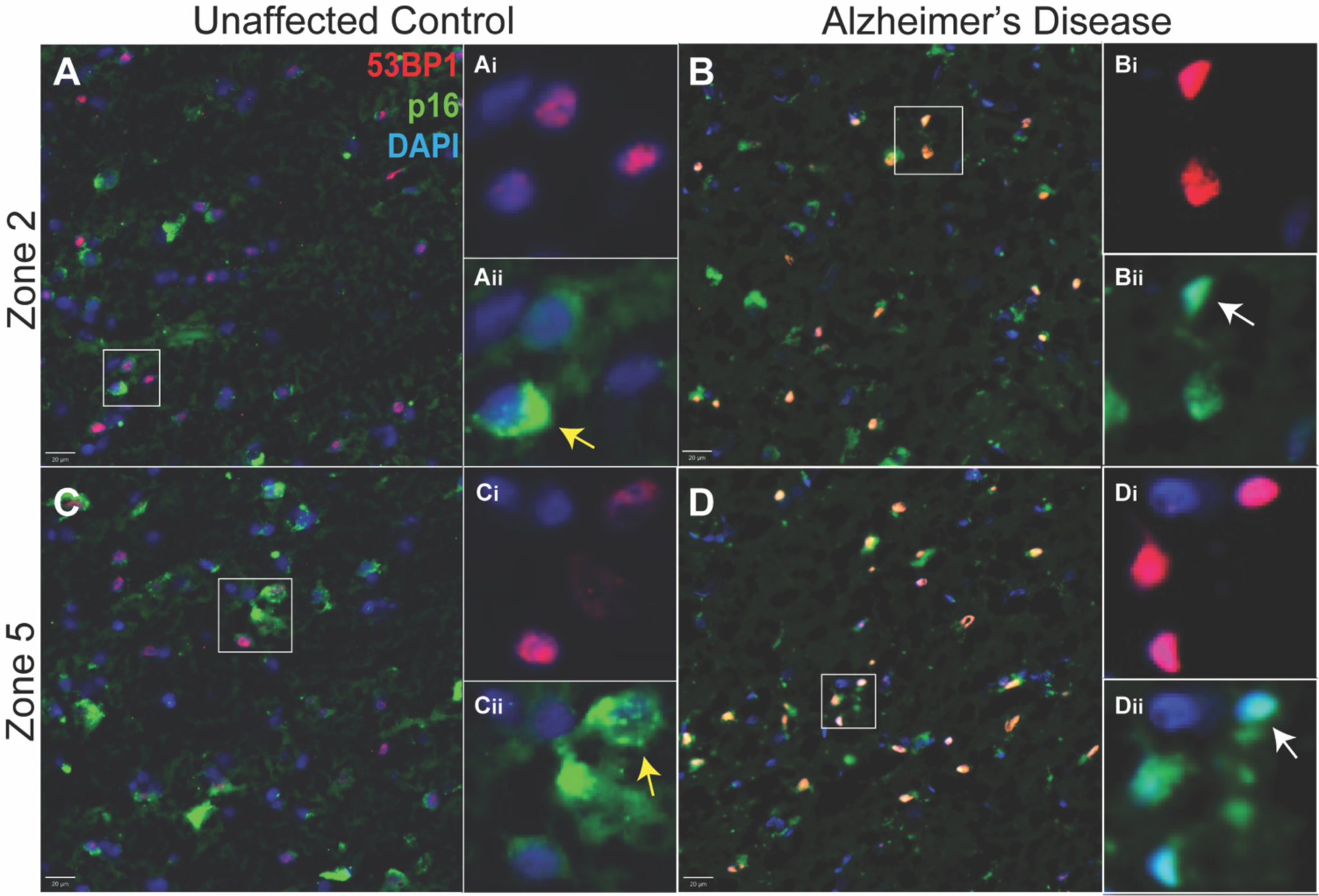
Examples of human brain cryostat sections immunostained for DNA damage (53BP1 – red) and senescence (p16 – green) with a DAPI (blue) nuclear counterstain. The panels on the left (**A, C**) are from an individual who died with no cognitive impairment. The panels on the right (**B, D**) are from a person who died with advanced Alzheimer’s dementia. Different cortical depths are illustrated for both cases. The top panels (**A, B**) are from a more superficial location (Zone 2); the bottom panels (**C, D**) are from a location closer to the white matter (Zone 5). The insets in each figure (e.g., Ai and Aii) are higher magnifications of the cells indicated by the white box.

3.2. *DNA damage increases with cortical depth in AD but not UC*

We next asked whether the density of cells displaying evidence of DNA damage or senescence changed in any consistent way with cortical depth. We performed a large, high-resolution scan of the immunostained human tissue then imported the image into QuPath ^30^. We divided the cortical gray matter into six radial zones as described in the Methods (Fig. 4A). We used DAPI staining to train the software to identify cell nuclei, then set the QuPath detection parameters in the red channel to pick only the more brightly labeled 53BP1-positive nuclei (Fig.4B). A representative field with the QuPath detections visible illustrates the outcome (Fig. 4C). When we did the same procedure for the AD cases, the nuclear intensity of 53BP1 staining and the number of cells quantified as being 53BP1+ were visibly increased (Fig. 4D and E). The density of total cells (DAPI+ nuclei) in the annotated area was not significantly different between AD and UC cases (Fig. 4F), although there was a slight trend towards increased density in the AD samples, possibly reflecting the atrophy of the neuropil. We then tallied the number of 53BP1+ cells in the AD cases and compared the values with those in the UC cases. We found a significant increase in the fraction of cells with DNA damage in the AD cases (Fig. 4G). These data confirm that cells with DNA damage are present in the aging human brain and that their density increases in Alzheimer’s disease.

**Figure 4.**
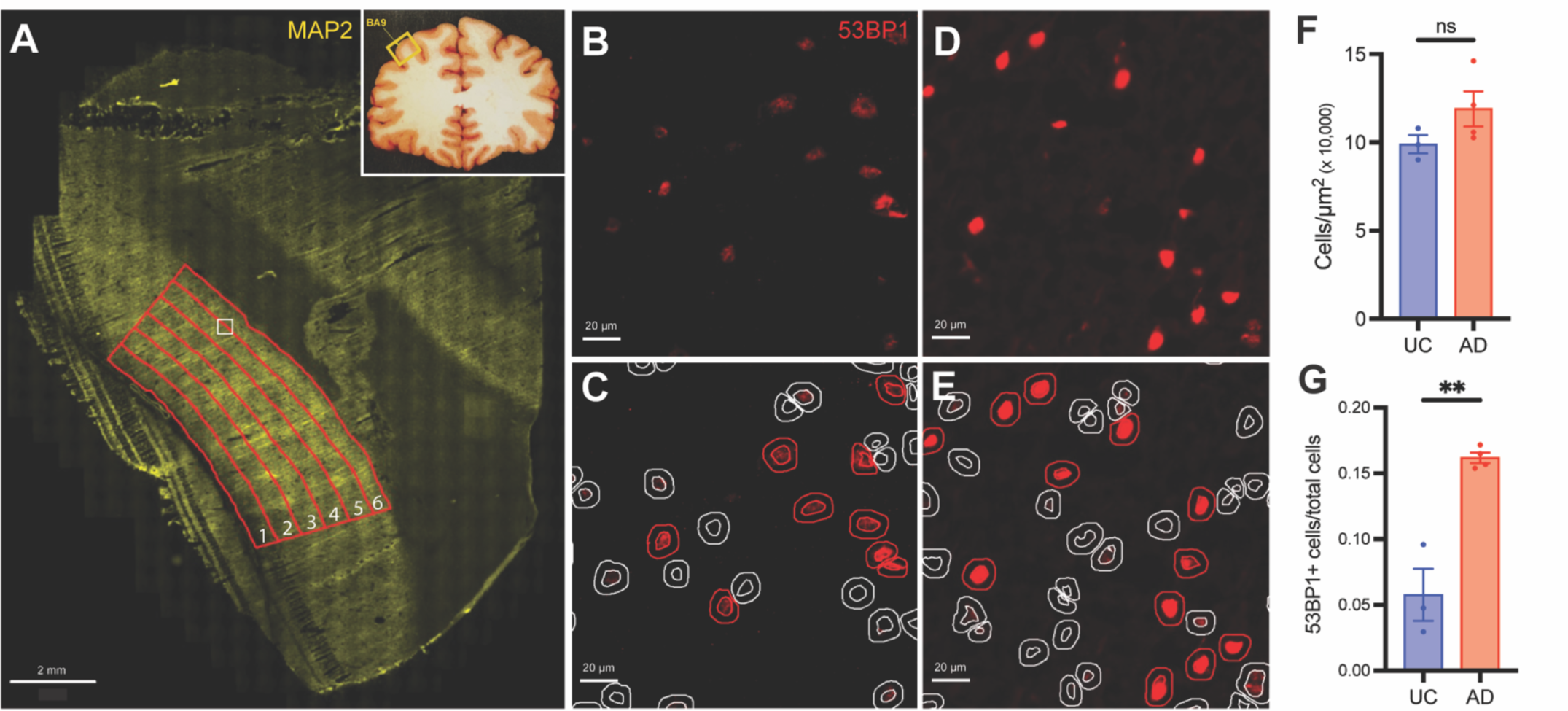
**A**) A low magnification image of a MAP2 immunostained section from human UC (Case 5794) BA9 (location indicated in the inset diagram). The red lines illustrate the six cortical zones. The leftmost zone that is closest to the pia is defined as zone 1 and the rightmost zone adjacent to the white matter is defined as zone 6. **B**) A high magnification image from the area indicated by the white box in panel **A** immunostained for 53BP1. **C**) The field of cells shown in panel **B** overlaid with the detections identified by QuPath. Inner circles represent the location of the DAPI stained nuclei (not shown); outer circles represent the “pseudoplasm” calculated as a 5 µm extension of the nuclear outline. Red detections represent cells identified as 53BP1-positive; grey detections represent 53BP1-negative cells. **D)** A high magnification image from Zone 5 in age-matched Human AD case stained with 53BP1. **E)** The field of cells shown in panel **D** overlaid with the detections identified by same QuPath parameters as panel **C**. **F**) Average density of DAPI^+^ nuclei in all AD or UC cases. **G**) Average density of 53BP1^+^ nuclei in all AD or UC cases. Error bars in **F** and **G** = SEM. AD: n=4 cases. UC: n=3 cases. (ns = not significant; ** p < 0.01).

In the UC cases, approximately 5-6% of the cells were 53BP1-positive and the density of these damaged cells changed little with cortical depth (Fig. 5A, B). We performed a least-squares-fit analysis of the data and while the line so defined trended higher with depth, the slope was not significantly different from zero (p = 0.15). This suggests a uniform distribution of DNA damage across the gray matter in the UC brains. In the AD brain, the density of 53BP1+ cells was significantly higher than in UC (by two-way ANOVA) with a different spatial distribution across the cortical depth. From the cortical surface (Zone 1) to the region we have called Zone 4 the density of cells increased steadily. In the deepest layers of cortex (Zone 6), however, the density of cells with DNA damaged decreased from Zone 5. Indeed, we found that a quadratic equation (r^2^ = 0.71, adjusted r^2^ = 0.68) produced a much better fit than a simple linear regression (r^2^ = 0.40) for the AD cases across the cortical depth, suggesting the curve is distinctly curvilinear (Fig. 5A, C). This idea is supported by the finding that the density of 53BP1^+^ cells in only Zones 3-5 was significantly higher than in Zone 1 (p < 0.001 for Zones 4 and 5). Compare this with the UC data, which is nearly identically fit with either the linear (r^2^ = 0.124) or the quadratic (r^2^ = 0.155, adjusted r^2^ = 0.043) regression. Thus, DNA damage follows an inverted U-shape trend across the cortical depth in the AD cases rather than the more linear trend seen in the UC cases. Together, these findings suggest not only that DNA damage is increased in AD cortex, but this AD-related increase is greater in the lower layers of the cortex compared to the upper layers.

**Figure 5.**
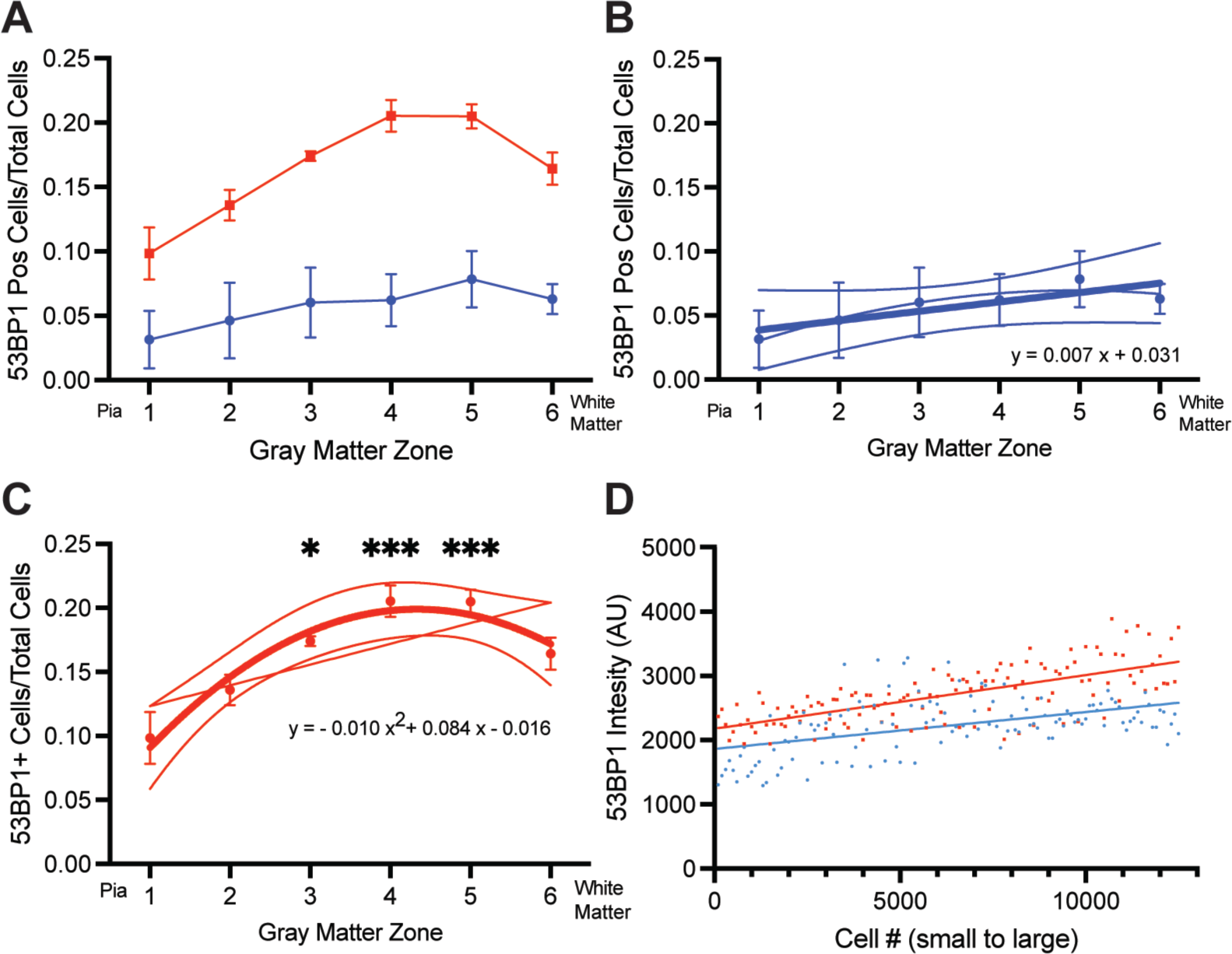
**A**) Average density of 53BP1^+^ cells in the six cortical zones of the UC (blue symbols) and AD (red symbols) cases. Error bars = SEM. AD: n=4 cases. UC n=3 cases**. B**) UC data from **A** replotted to show the least squares fit relationship (thick line, r^2^= 0.124) and the quadratic fit (thin line, r^2^ = 0.155, adjusted r^2^ = 0.043) with 95% confidence limits. The least squares method gave a slightly better fit to the data, but the slope was not significantly different from zero (p= 0.1522). **C**) AD data from **A** replotted to show the least squares fit relationship (thin line, r^2^= 0.40) and the quadratic fit (thick line, r^2^= 0.71) with 95% confidence limits. The quadratic equation gave a much better fit to the data. **D**). A representative plot of 53BP1 intensity as a function of nuclear size. The X-axis represents the ranking of over 12,000 cells from Zone 4 by the area of their nuclei (from smallest to largest). Each point represents the average of 100 cells. The Y-axis is the average intensity of 53BP1 staining in successive groups of 100.

The relationship between DNA damage and cortical depth was not a reflection of the prevalence of amyloid plaques. Although the Aβ peptide has been cited as an inducer of DNA damage ^31^ the pattern of amyloid deposition does not match pattern of DNA damage (Fig. 1 and 5) suggesting that amyloid induced DNA damage is not the underlying cause of the pattern shown in Figure 1C. Our analysis does not differentiate between neuronal and non-neuronal cells, as we did not include a cell type marker in our large scans. In separately stained sections (e.g., Fig 2), we regularly observed evidence of DNA damage in neurons and non-neurons alike. To test whether this was consistent with the findings from the large scan, we sorted the cells in both AD and UC cases in order of increasing nuclear size. The nuclei of neurons are significantly larger than non-neurons. Therefore, if neurons have more DNA damage than non-neurons, the intensity of *γ*-H2AX immunostaining should be more intense in cells with large nuclei. The representative graph in Figure 5D illustrates that there is a slight trend in this direction; the slopes of both the AD and UC lines are significantly different from zero (p < 0.001), but the shallow slopes suggest that the relationship is not as strong as would be expected if cell type were a significant factor in the extent of DNA damage. Note as well that the differences between AD and UC are apparent at every cell size from the smallest to the largest. The suggestion that emerges is that all cell types are affected by age and by Alzheimer’s disease.

### 3.3. The p16^INK4a^ senescence marker is found in both nucleus and cytoplasm

We also found evidence for the presence of senescent cells in the aging and Alzheimer’s disease brain using p16^INK4a^ as a marker (Fig. 3). The analysis of the senescent phenotype was made more complicated because p16 staining appeared in two distinct patterns. The first was a clear, though sometimes diffuse, nuclear presence (white arrows, Figs. 3Bii and 3Dii). The second was a clumped, often punctate, cytoplasmic appearance (yellow arrows, Figs. 3Aii and 3Cii). Nuclear and cytoplasmic patterns were found in both AD and UC cases. Most cells tended to have either one pattern or the other, but there were clearly cells that displayed p16 in both locations. This complex situation made it difficult to accurately count a simple metric such as the number of p16-positive cells.

Beginning with the signal only in the nucleus, we found that the average nuclear intensity of p16 was greater in AD brains than in UC (Fig. 6A). We then focused solely on the cells identified with DNA damage (53BP1+) and quantified the intensity of p16 in cell nuclei and cell cytoplasm separately. Plotting p16 nuclear intensity as a function of cortical depth (Fig. 6B) revealed a weak, but significant trend towards increased intensity in the lower layers of cortex. The same nearly linear relationship was seen for both UC and AD with both slopes significantly different from zero (p = 0.0377 and p < 0.0001 respectively). A two-way analysis of variance (ANOVA) revealed a significant AD vs UC effect (p= 0.0002) and a significant zone effect (p= 0.0046) with a nonsignificant interaction (p= 0.9978) on average nuclear intensity. To measure cytoplasmic staining, we used the QuPath “cell expansion” feature, to draw a concentric line at a set distance (5 µm) from the outline of the DAPI-defined nuclear boundary. We refer to the area between the nuclear and expansion as the cytoplasm but recognize that it is only a sample of the full cytoplasm. As with nuclear p16, cytoplasmic p16 staining also showed a modest tendency to increase with cortical depth (Fig. 6C). The slopes of these lines were reduced from those in Fig. 6B, and only the AD curve was significantly different from zero. Further, unlike the nuclear staining, there was no difference between the staining intensity of the cytoplasmic p16 in AD compared with UC with a nonsignificant AD vs UC effect (p= 0.1828).

**Figure 6.**
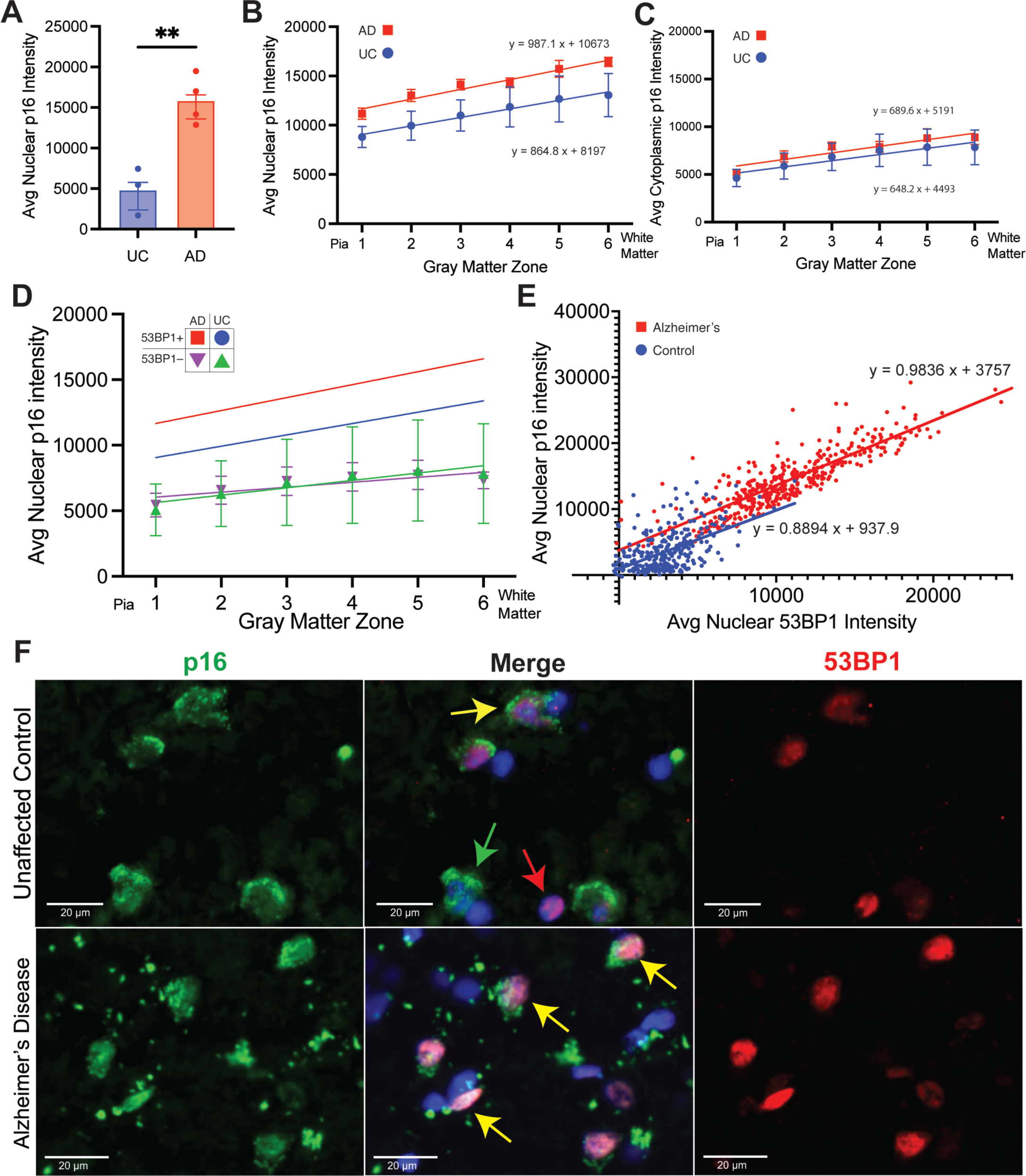
**A**) Average intensity of p16 immunostaining in the nuclei of cells identified as 53BP1^+^ in both UC (blue) and AD (red) cases. ** p < 0.01. **B**) Average intensity of nuclear p16 immunostaining of 53BP1^+^ cells in the six cortical zones of the UC (blue symbols, r^2^= 0.2429 with significant non-zero slope (p=0.0377))) and AD (red symbols, r^2^= 0.7024 with significant non-zero slope (p<0.0001))) cases. **C**) Average intensity of “pseudoplasmic” p16 immunostaining of 53BP1^+^ cells in the six cortical zones of the UC (blue symbols, r^2^= 0.1958 with nonsignificant non-zero slope (p>0.05))) and AD (red symbols, r^2^= 0.4843 with significant non-zero slope (p=0.0002))) cases. **D**) Average intensity of nuclear p16 immunostaining of 53BP1^+^-negative cells in the six cortical zones of the UC (green symbols) and AD (purple symbols) cases. Error bars in **A-D** = SEM. AD: n=4 cases. UC: n=3 cases. **E**) A plot of nuclear p16 immunostaining intensity as a function of 53BP1 immunostaining intensity from a random sampling of 100 cells/case from UC (blue symbols, n=3) and AD (red symbols, n=4) cases. The equations for the least squares fit of the two data sets are indicated. **F:** Immunohistochemistry showing the relationship and colocalization of p16 and 53BP1 in the nucleus. Green arrows indicate cells positive for p16 only. Red arrows indicate cells positive for 53BP1 only. Yellow arrows indicate cells positive for both p16 and 53BP1.

The data suggest a robust relationship between cells with DNA damage and cells with evidence of senescence. To emphasize this point, we performed an identical analysis using only cells that we had scored as 53BP1-negative (purple and green symbols, Fig. 6D). This data revealed two important features. The first was that, independent of the UC or AD source of the tissue (nonsignificant AD vs UC effect, p= 0.9448), the p16 immunostaining was much weaker in cells with no overt evidence of DNA damage. The second was that in the UC, even the weak staining that was present in the DNA-damage-negative cells showed little or no evidence of change with depth in the cortex. The AD values did increase slightly but the r^2^ value of 0.2352 suggests that the relationship was not strong. Finally, we asked whether the relationship between DNA damage and senescence existed on a cell-by-cell basis. We plotted the intensity of nuclear p16 staining as a function of 53BP1 intensity (Fig. 6E). Most cells in the UC cases formed a cloud of points near the origin of the graph with a moderate positive linear relationship between the two variables (blue symbols, Fig. 6E; r^2^ = 0.2805). In the AD cases by contrast, there was a strong linear relationship between the level of DNA damage and senescence (red symbols, Fig. 6E; r^2^ = 0.7210). We noted as well that for the few cells in the UC cases where p16 was more intense, their levels of 53BP1 staining placed their position on the graph close to that expected from a cell in an AD case.

The staining pattern itself suggested another perspective on the damage/senescence relationship. In the UC, cells with nuclear p16 staining were often negative for 53BP1 (Fig. 6F, top row, green arrows) and there were cells with evidence of DNA damage, but none for senescence (Fig. 6F, top row, red arrow). There were also cells with both p16 and 53BP1 evident (Fig. 6F, top row, yellow arrow), but in general the staining intensity of markers in these cells was weak as would be predicted from the cluster of blue points near the origin in Figure 6E. In the AD brain, by contrast, there was a much greater tendency for the two markers to overlap (Fig. 6F, bottom row, yellow arrows). The strengthening of the signal coupled with the strong overlap, can even be seen at lower magnifications (Fig. 3B, D).

The strong correlation of DNA damage and senescence in the Alzheimer’s disease brain suggested that there could be a causal relationship between the two cellular phenomena. We hypothesized, based on the literature, that the most likely nature of this relationship was that DNA damage induced senescence. To test this idea, we turned to in vitro culture of mouse embryonic cortical neurons. Neuron-enriched cultures were plated on coverslips in 24-well plates and allowed to mature for 14 days in vitro (DIV14). We then induced DNA damage with etoposide (ETOP), a drug that inhibits the activity of topoisomerase-II. Twenty-four hours after drug treatment, we fixed the cells and immunostained for MAP2 (a neuronal marker), *γ*-H2AX to reveal DNA damage and p27 (a second marker of cellular senescence). The health of the cultures in general, and the integrity of the neuronal processes in particular, were unaffected by the treatments applied (Fig. 7A, MAP2 staining in left column). Nonetheless, 10 µM etoposide had its intended effect; after 24 hours the percentage of cells with DNA damage (*γ*-H2AX staining) increased from the background levels of untreated cells to nearly 40% (Fig. 7B). Despite this dramatic increase in DNA damage, however, the fraction of senescent cells remained unchanged (Fig. 7C). The absence of a senescence response can also be seen on a cell-by-cell basis. For the control cultures, the graph of p27 intensity as a function of DNA damage (*γ*-H2AX immunostaining) is basically a cluster of points around the origin (Fig. 7D) with a group of cells that have no DNA damage, but significantly increased p27 expression. For the cultures treated with 10 µM etoposide (Fig. 7E), the extent of DNA damage increased. Despite this, the levels of p27 showed no tendency to increase within the cells with greater *γ*-H2AX staining.

**Figure 7.**
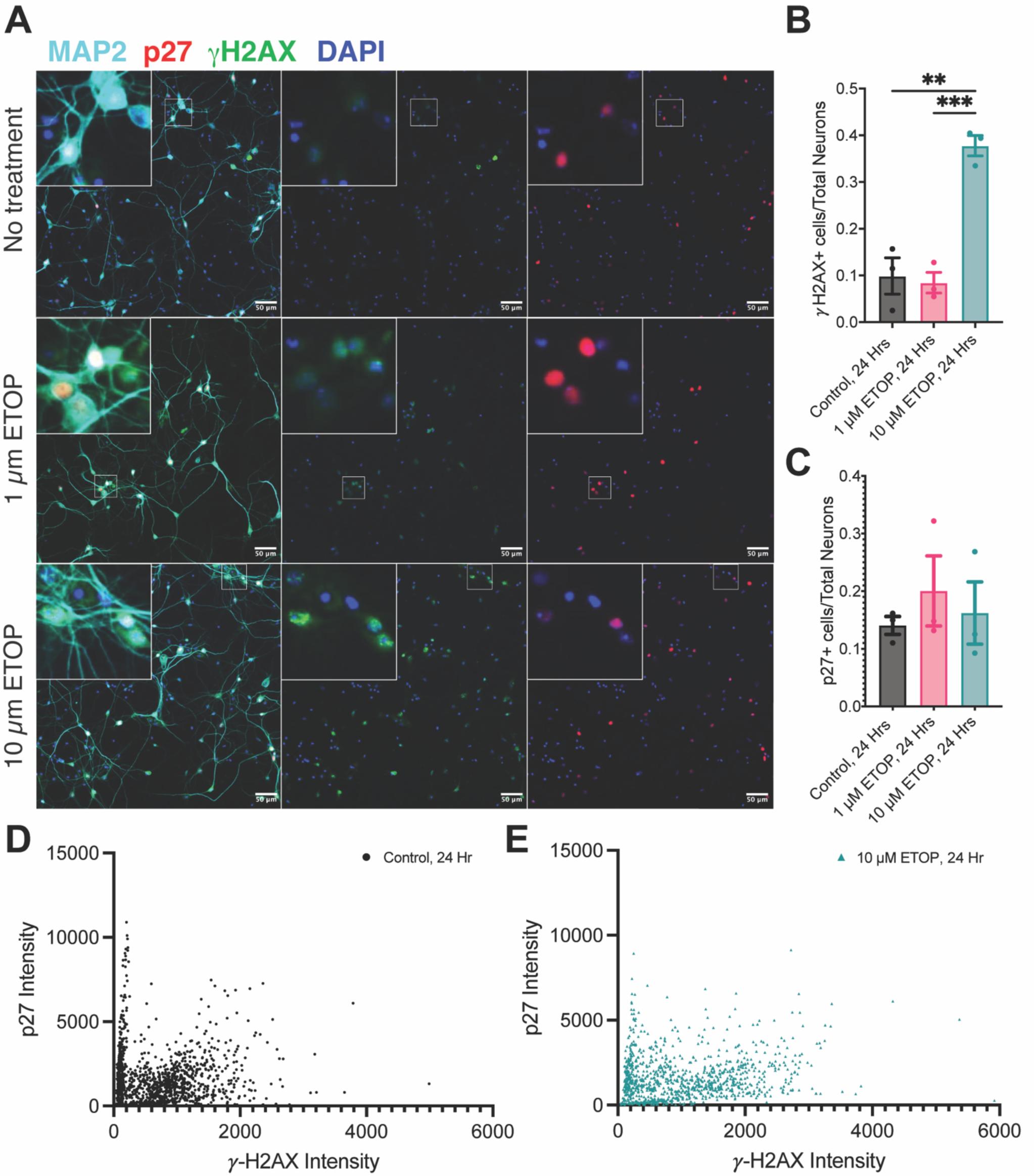
**A**) Representative fields of DIV14 cultures of mouse embryonic cortical neurons immunostained for MAP2 (cyan), p27 (red), *γ*-H2AX (green) and counterstained with DAPI (blue). The top row of images are from a control culture (n=3); the middle row is from cultures treated for 24 hr with 1 µM etoposide (n=3); the bottom row is from cultures treated with 10 µM etoposide (n=3). Left column = merge; middle column = *γ*-H2AX plus DAPI; right column = p27 plus DAPI. **B**) The fraction of *γ*-H2AX-positive cells in control and etoposide treated cultures (** p < 0.01; *** p < 0.001). **C**) The fraction of p27-positive cells in control and etoposide treated cultures. Error bars in **B** and **C** = SEM. **D**) A plot of p27 immunostaining intensity as a function of *γ*-H2AX immunostaining intensity of cells in untreated cultures (n=3). **E**) p27 immunostaining intensity as a function of *γ*-H2AX immunostaining intensity of cells in cultures treated with 10 µM etoposide for 24 hr (n=3).

## 4. Discussion

Damage to the cellular genome is tightly correlated with the process of aging ^11, 32^ and has been described as a driver of aging rather than merely a symptom ^3, 5, 6, 8, 15^. Our data confirm that evidence of DNA damage is present in the cells of the aging brain. In prefrontal cortex, Brodmann Area 9 (BA9), we find that approximately 5% of the cells are immunopositive for 53BP1, part of a complex of proteins that assembles at sites of DNA double strand breaks, and for *γ*-H2AX, a modified histone that is found in the regions surrounding regions of DNA repair. In samples from the BA9 region of persons who died with Alzheimer’s disease, the fraction of neurons with DNA damage approached 20% – 3 to 4-fold higher than controls. While this result may have been predictable given the literature on the topic, our data offers further insight by bringing anatomical resolution to the data. In the unaffected control samples we examined, the small number of cells with DNA damage tended to scatter uniformly across the cortical layers. In the AD brain, we observed an increase of DNA damage as the cortical layers become deeper. However, our quadratic regression analysis showed the increase is not uniform. Rather, in the more superficial Zone 1 the density of cells that were positive for 53BP1 staining was relatively low (although still higher than UC). With increasing cortical depth, the density increased to a maximum in Zones 4 and 5 followed by a clear decrease in the deepest zone, Zone 6. In addition, from the quadratic regression curve, our mathematical model predicts that between Zone 4 and Zone 5 is where the most DNA damage is to be found. The cortical zones as we have defined them are not identical with the histologically defined cortical layers, but we may use them as reasonable surrogates. This suggests that the cells of Layers III, IV and V are the most vulnerable to damage during the progression of AD. Layers IV and V are the major input and output regions of cortex respectively.

It has been noted that increased neuronal activity induces transient DNA damage ^20, 24, 25^. With age and the onset of Alzheimer’s disease, however, DNA breaks become more prevalent and their repair less efficient. From this vantage point one interpretation of our findings would be that neuronal activity is highest in these regions, and in the AD brain the counteracting repair mechanisms are overwhelmed leading to a state of persistent damage. This idea is consistent with the observation of increased epileptic activity in human AD and its mouse models ^33, 34^. Other explanations are possible, but we propose that the data are incompatible with a model in which the accumulation of DNA damage is a random or haphazard event whose probability is equivalent in all regions and all cells. The pattern of DNA damage in our material also matches the density of tangles of hyperphosphorylated tau in the BA9 region of AD brains ^35^ while there is a poor match with pattern of plaques Fig. 1 plus images in reference ^36^. As the density of tau deposits is well correlated with the extent of neurodegeneration, the connection among the patters of neurofibrillary tangles, DNA damage, and senescence takes on added meaning. Notably, it has also been observed that other pathological changes in Alzheimer’s disease that are not correlated with the distribution pattern of plaques such as the axon initial segment^37^.

The strength of the correlation between senescence and DNA damage adds another important dimension to our findings. The sub-cellular distribution of the senescence marker, p16 (Fig. 1) is unexpected but not unprecedented. Other laboratories have reported on the presence of p16^INK4a^ in the neurons of the aging brain and its increased presence in Alzheimer’s disease ^38-40^. The immunostaining pattern in these early reports matches our immunofluorescent staining quite well, with both nuclear and cytoplasmic staining visible in the published figures. The relationship of p16 and 53BP1 staining is notable as DNA damage has been identified as a potential trigger of cellular senescence in dividing cells and cell lines ^29^. Less is known, however, about the impact of DNA damage with regard to the senescence process in highly differentiated cells such as those in the brain. While there is clearly a strong correlation between the two, the data in the UC and AD cases illustrated in Figure 6E contain the intriguing suggestion that the relationship may not be a simple go/no-go decision. Rather senescence responds in an analog rather than a digital fashion to DNA damage. Further, while we recognize that primary cultured neurons are not a fully representative model of the complex phenotype of cells in the aging human brain, direct production of DNA damage with etoposide does not induce a senescent phenotype in cultured neurons within the 24 hour period of our experiment. It is possible that longer incubation times following the etoposide challenge might be needed to fully test the lack of a causal relationship. The results nonetheless raise the possibility that while damaged DNA might be a necessary part of the program of cellular senescence, it may not be sufficient on its own. The increased overlap of DNA damage and senescence seen in the AD but not the UC brain suggests that there is shift that occurs in the chemistry of the AD brain and this shift has effects on both the level of DNA damage and the senescent response. The nature of this shift is unknown.

The results sharpen our understanding of the interplay of DNA damage and senescence in the aging brain, but in doing so, they raise new questions to be answered in future studies. What is the relative timing of the appearance of genomic damage and the senescent phenotype. What is it about the chemistry of the AD brain that triggers both senescence and DNA damage? Our study was restricted to cases of sporadic AD. Do the mouse models of AD, which reproduce familial AD, show the same phenotypes? The answers are important as both senescence and loss of genomic integrity threaten the health and proper function of the adult brain and factors leading to the loss of either or both may hold important clues for the causes and treatments of dementing illnesses such as Alzheimer’s disease.

